# Constitutive Immune Activity Promotes Tumorigenesis in *Drosophila* Intestinal Progenitor Cells

**DOI:** 10.1101/085597

**Authors:** Kristina Petkau, Meghan Ferguson, Silvia Guntermann, Edan Foley

## Abstract

Gut innate immune defenses contain bacterial populations and protect the host interior from invasive microbes. Although excess intestinal immune activity frequently promotes inflammatory illnesses, we know little about the consequences of chronic innate immune activity exclusively in endodermal gut cells of an otherwise normal animal. To address this question, we generated a transgenic line that allows us to activate inflammatory signals in adult fly intestinal progenitor cells. We found that constitutive immune activity in intestinal progenitors disrupts expression of homeostatic regulators such as Notch signal transduction pathway components and induces hyperplasia throughout the gut. Consistent with these observations, we found that persistent immune signaling interferes with progenitor cell differentiation and exacerbates the formation of Notch-dependent intestinal tumors. These findings uncover a novel link between constitutive immune activity and tumorigenesis in intestinal stem cells.

## INTRODUCTION

The endoderm is at the heart of an ancient, intimate, and continuous relationship between multicellular organisms and the microbial world. Gut commensals negotiate a semi-stable existence within host niches, partially through provision of factors that influence nutrition, development, and immunity in the host (Cho and Blaser, 2012). For their part, hosts invest energy and resources in the containment of intestinal microbes (Hooper et al., 2012). Physical barriers such as the peritrophic matrix of insects, or dense mucosal layers of mammals, keep microbes at an adequate distance from gut cells. Additionally, host-derived reactive oxygen and nitrogen species destroy invading microbes, while antimicrobial effectors block microbial dissemination. The consequences of failed antibacterial defenses are often extreme, and occasionally deadly for the host. For example, dysbiotic microbial communities contribute to traumatic inflammatory bowel diseases in humans (Wlodarska et al., 2015). In recent years, we have made considerable advances in understanding relationships between host immune responses and gut microbiota (Honda and Littman, 2016). Germline-encoded innate immune systems provide critical defenses in the endoderm, a primary site of contact between multicellular organisms and their immediate microbial environment. Thus, characterization of innate immune regulation in endodermal tissues is essential for a full appreciation of the mechanistic basis for the restraint of gut microbes.

The fruit fly, *Drosophila melanogaster,* is widely used for the characterization of intestinal development and function (Buchon et al., 2013, Jiang and Edgar, 2012, Lemaitre and Miguel-Aliaga, 2013). For example, *Drosophila* is an excellent model for the exploration of Notch-mediated regulation of gut development (Guo and Ohlstein, 2015, Micchelli and Perrimon, 2006, Ohlstein and Spradling, 2006, Ohlstein and Spradling, 2007), and intestinal tumorigenesis (Patel and Edgar, 2014, Biteau and Jasper, 2011, Marianes and Spradling, 2013, Apidianakis et al., 2009). In the fly midgut, basal intestinal stem cells (ISC) divide to generate a bipotent transient cell type, the enteroblast (EB). Delta-Notch signals between ISC-EB progenitor pairs determine the developmental fate of EBs (Takashima et al., 2011, Perdigoto et al., 2011). High levels of Notch activity in enteroblasts lead to their differentiation as large, polyploid, absorptive enterocytes (EC). Lower levels of Notch activity result in the differentiation of enteroblasts as smaller, diploid, secretory enteroendocrine cells (EE). Notch-dependent control of progenitor cell differentiation is conserved across vast evolutionary distances, with similar requirements in species as diverse as fish and rodents (Fre et al., 2005, Stanger et al., 2005, van Es et al., 2005, Crosnier et al., 2005), and interruptions to Notch signaling lead to intestinal tumor formation in several models (Kazanjian and Shroyer, 2011, Peignon et al., 2011).

Current data implicate the immune deficiency (IMD) pathway as a central regulator of antibacterial defenses in the fly gut (Buchon et al., 2009, Lhocine et al., 2008). IMD shares numerous features with the mammalian tumor necrosis factor (TNF) cascade (Buchon et al., 2014). In the fly, detection of bacterial peptidoglycan results in the proteolytic removal of thirty N-terminal amino acids from the Imd adaptor protein by the caspase Dredd (Paquette et al., 2010). Cleaved Imd associates with the Inhibitor of Apoptosis 2 (IAP2) and Fas-Associated Death Domain (FADD) orthologs (Guntermann and Foley, 2011, Paquette et al., 2010) to activate the NF-kB transcription factor Relish (Dushay et al., 1996). Active Relish relocates to the nucleus and initiates transcription of a broad spectrum of antimicrobial peptides. More recent work showed that intestinal IMD activity also controls the expression of genes involved in developmental and metabolic processes (Broderick et al., 2014, Erkosar et al., 2014). These studies hint at novel, unstudied biological roles for IMD that extend beyond the extermination of unwanted bacteria.

Here, we characterized the consequences of persistent IMD activity in midgut progenitors. Midgut progenitor cells extend projections into the intestinal lumen (Ohlstein and Spradling, 2006), and express IMD pathway genes at levels comparable to mature epithelial cells (Dutta et al., 2015). In addition, enteric challenges with bacterial pathogens induce classic IMD-response genes in progenitor cells (Dutta et al., 2015, Fink et al., 2016), confirming progenitor cells as an immune-competent compartment. However, the effects of IMD activity on progenitor cells is unclear. To address this, we engineered a novel IMD variant that permits temporal and cell-specific activation of IMD. We showed that constitutive immune signaling altered the expression of Notch pathway regulators, disturbed EE differentiation, and caused hyperplasia throughout the posterior midgut. Consistent with interactions between IMD and Notch, we found that constitutive IMD activity greatly exacerbated the formation of Notch-dependent tumors. Given the involvement of inflammation and Notch signaling in vertebrate intestinal tumor development, we believe our findings may be relevant to studies that explore relationships between intestinal inflammation and tumorigenesis.

## RESULTS

### Constitutive IMD Pathway Activation in Intestinal Stem Cells

To characterize the effects of constitutive immune signaling in the intestine, we generated a transgenic *Drosophila* line that allows cell-restricted activation of the IMD pathway. Specifically, we engineered a truncated IMD protein that uses an internal ATG (at residue 78) as a start codon. We refer to this truncated protein as ImdCA. ImdCA lacks inhibitory N-terminal amino acids (Figure 1A), but retains the ability to interact with the FADD adaptor protein (Figure 1B). We generated transgenic fly lines that permit temperature-dependent expression of ImdCA in fat tissue (*cgGAL4* ; *GAL80^ts^*/*UASimdCA* (abbreviated as *cg^ts^>CA*)), or ISC/EB progenitor cell pairs (*esgGAL4, GAL80^ts^, UASGFP*; *UASimdCA* (abbreviated as *esg^ts^>CA*)). Incubation of either genotype at 29°C, the restrictive temperature for *GAL80^ts^*, results in tissue-specific expression of ImdCA in fat body or intestinal progenitor cells, respectively. In the fat body, the principle site of humoral immunity, ImdCA caused expression of the IMD-responsive antimicrobial peptides *dpt* and *att* (Figure 1C). The induction of *dpt* and *att* proceeded via a classical IMD response, as null mutations in *dredd* prevented ImdCA-mediated antimicrobial peptide expression (Figure 1D). ImdCA is equally effective at inducing antimicrobial expression in midgut progenitor cells (Figure 1E), and expression of ImdCA provided protective benefits against lethal challenges with the gastrointestinal pathogen *Vibrio cholerae* (Figure 1F). These data show that we have established a novel fly line that permits tissue-specific activation of IMD responses in *Drosophila.*

**Figure 1.**
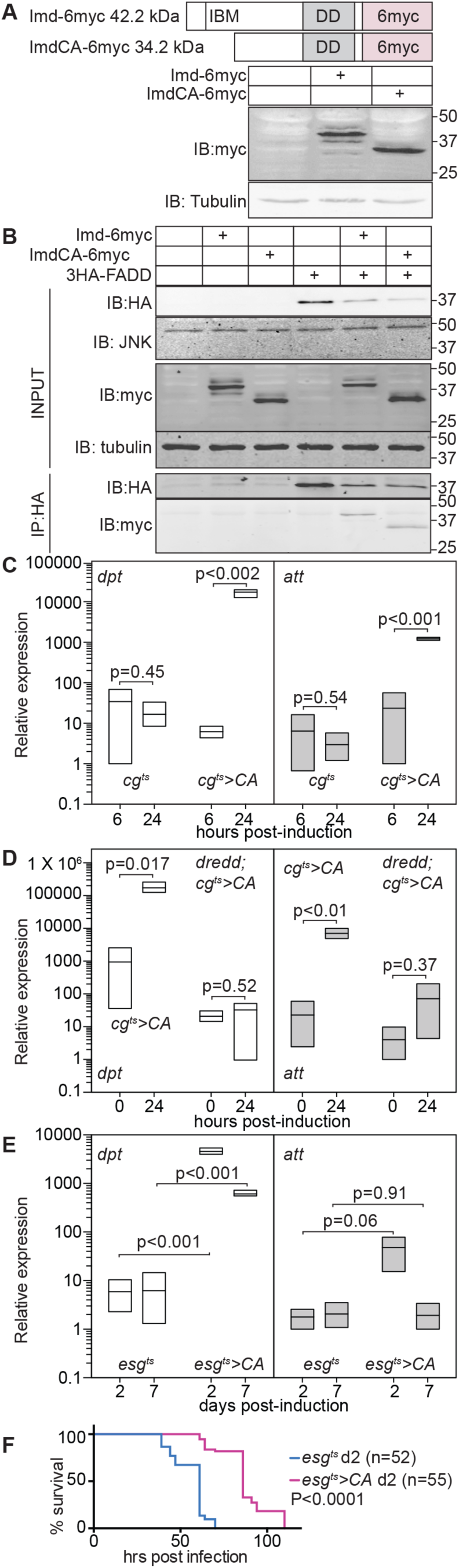
A truncated Imd protein activates innate immune responses. **(A)** Schematic representation of C-terminally myc-tagged Imd and ImdCA. Iap Binding Motif (IBM) and death domain (DD) are indicated. Western blots show lysates of untransfected S2 cells (lane 1) S2 cells transfected with expression constructs for Imd-6myc (lane 2), and ImdCA-6myc (lane 3). Numbers indicated molecular weights in kDa. The membrane was probed with Tubulin as a loading control. **(B)** S2 cells were transfected with the indicated expression constructs and input samples (blots labeled INPUT) were probed for the indicated antigens. Anti-HA immunoprecipitates (blots labeled IP:HA) were probed with the indicated antibodies. **(C-E)** Quantification of the relative expression levels of *dpt* and *att* in flies raised at 29°C for the indicated time. Survival curves of *esg^ts^>CA* and *esg^ts^* flies raised at 29°C for two days, and then challenged with *Vibrio cholerae.*

### A Balance of Microbial Cues and Host Immune Status Determines Gene Expression Patterns in the Adult Midgut

Intestinal progenitor cells express IMD pathway components at similar levels to epithelial cell (Figure 5A), and display classical IMD-responses to microbial challenges (Dutta et al., 2015). To characterize the effects of ImdCA expression in intestinal progenitors, we prepared transcriptional profiles of the intestines of *esg^ts^* and *esg^ts^>CA* adult flies that we raised at 18°C for ten days, and shifted to 29°C for two days. We examined *esg^ts^* and *esg^ts^>CA* flies raised under conventional or germ-free conditions in these studies. This strategy allowed us to make pairwise comparisons of the effects of commensal bacteria and ImdCA on the control of host transcription (Figure 2A-D).

**Figure 2.**
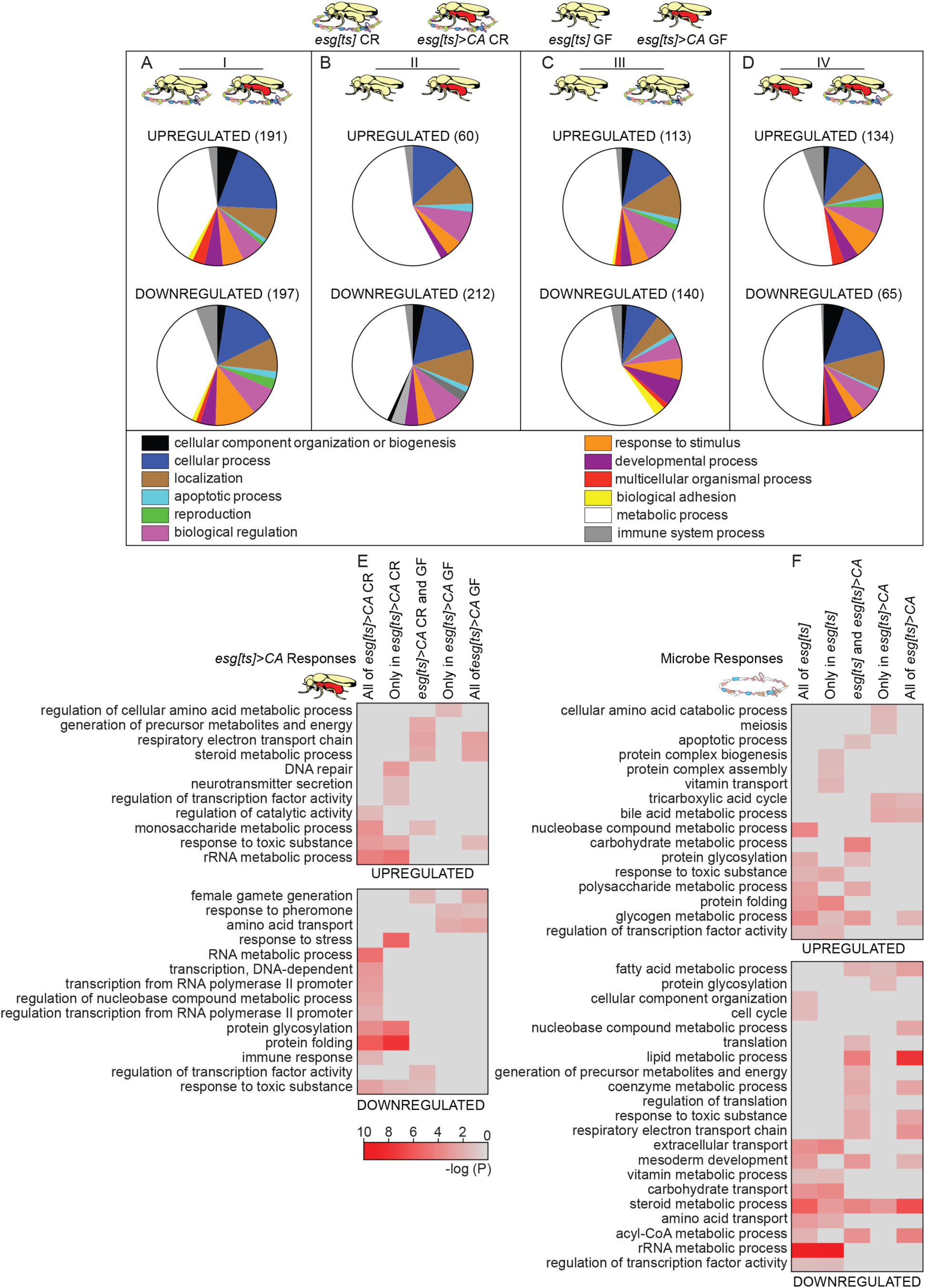
ImdCA and the microbiome interact to control gut transcription. **A-B:** Graphic representation of biological functions dysregulated in *esg^ts^>CA* flies relative to *esg^ts^* flies raised under conventional (A) or germ-free conditions (B). **C-D:** Graphic representation of biological functions dysregulated in conventional flies relative to germ-free flies in the absence (C) or presence (D) of ImdCA expression. For A-D, numbers indicated total numbers of dysregulated genes. Genes with unknown function were not included in pie charts. **(E-F)** Heatmap of significance scores for gene ontology terms (minimum 5 genes in each term) that were significantly dysregulated in *esg^ts^>CA* flies relative to *esg^ts^* flies (E), or conventionally-reared relative to germ-free flies (F). In E, GO terms are further subdivided into terms that were dysregulated in conventionally-reared or germ-free flies. In F, GO terms are subdivided into terms that were dysregulated in *esg^ts^* flies and *esg^ts^>CA* flies.

Similar to earlier studies (Broderick et al., 2014, Erkosar et al., 2014), we found that the microbiome regulates the expression a large cohort of host genes, with a particularly notable effect on genes involved in metabolic processes (Figure 2A-D, Figure 3). As expected, ImdCA altered the expression of genes with known roles in innate defenses. However, we also noticed effects of ImdCA on pathways as diverse as metabolism, transportation and stress responses (Figures 2, 3).

**Figure 3:**
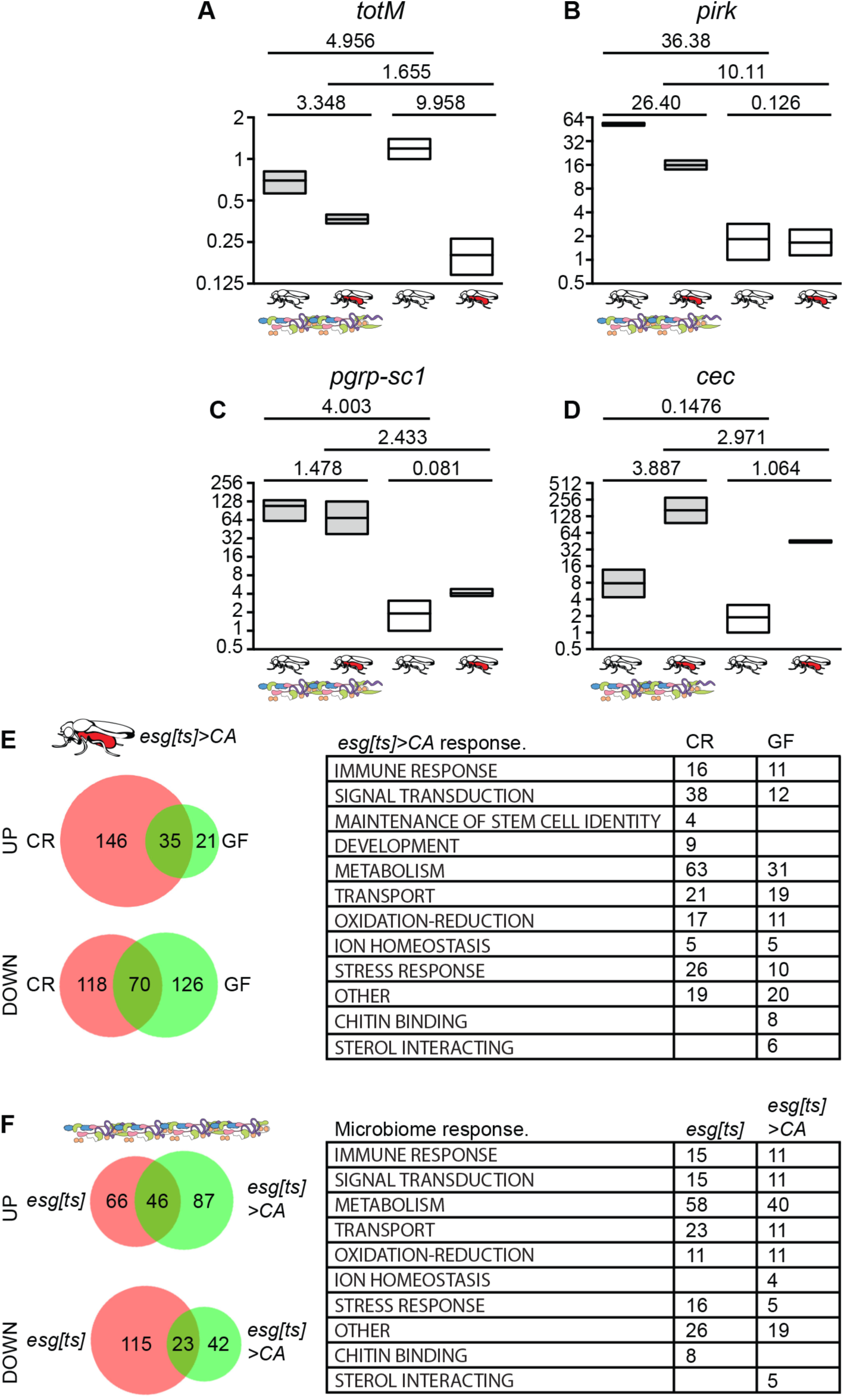
**A-D:** Relative expression of totM (A), pirk (B), pgrp-sc1 (C) and cec (D) in the dissected intestines of conventionally-reared or germ-free *esg^ts^* flies or *esg^ts^>CA* flies. For each assay, expression levels were reported relative to those observed in germ-free *esg^ts^* flies. Comparisons were performed with a Sidak’s multiple comparison test. Values show the Sidak’s t value for the respective tests. **E:** Number of genes and GO terms that were up or downregulated in *esg^ts^>CA* flies relative to *esg^ts^* flies raised under conventional (CR) or germ-free (GF) conditions. **F:** Number of genes and GO terms that were up or downregulated in conventionally-reared flies relative to germ-free flies of the indicated genotypes.

Comparisons between the individual expression profiles uncovered substantial effects of the microbiome on ImdCA-regulated transcription. For example, only 35 host genes are induced in both conventionally-reared and germ-free animals in response to ImdCA (Figure 3E). In contrast, 146 transcripts are only induced in the presence of an intact microbiome, and an additional 21 transcripts are only induced in the absence of a microbiome (Figure 3E). Likewise, microbiome-response genes are heavily influenced by the extent of immune activity in the gut. For example, a unique set of 129 genes responds to presence of the microbiome only upon induction of ImdCA (Figure 3F). GO term analysis showed that the microbiome and immune activity synergistically influence functional outputs in the gut. For example, ImdCA only promotes protein glycosylation in the presence of a microbiome (Figure 2E), and the gut microbiome induces tricarboxylic acid cycle genes only when accompanied by intestinal inflammation (Figure 2F). Combined, these data show that gut transcriptional outputs reflect the extent of immune activity and microbial presence in the gut.

Given the effects of ImdCA on antimicrobial peptide expression, we assumed that ImdCA will affect composition of the host microbiome. To test this, we treated freshly eclosed *esg^ts^* and *esg^ts^>CA* flies with a regime of antibiotics for six days to eliminate the endogenous microbiome. We then fed all flies a homogenate prepared from our lab wild-type *Drosophila* strain, and passaged flies to fresh food twice over seven days to facilitate stable association between microbes in the homogenate and the recipient flies. After seven days, we shifted recipients to 29°C for an additional seven days to activate the IMD pathway in *esg^ts^>CA* flies. We dissected the intestines of the different populations and used deep-sequencing of 16S DNA to identify the bacterial communities present in the respective guts (Figure 4). As expected, the bacterial microbiome of *esg^ts^* flies was barely distinguishable from the homogenate. Both microbiomes contained a limited number of OTUs characterized primarily by *Acetobacter* and *Lactobacilli* (Figure 4). Contrary to our initial expectation, we did not observe a significant impact of ImdCA on microbiome composition. In fact, the bacterial communities of the original homogenate, the *esg^ts^* flies, and the *esg^ts^>CA* flies were barely distinguishable (Figure 4), suggesting that acute activation of IMD does not significantly alter microbial populations in the gut.

**Figure 4.**
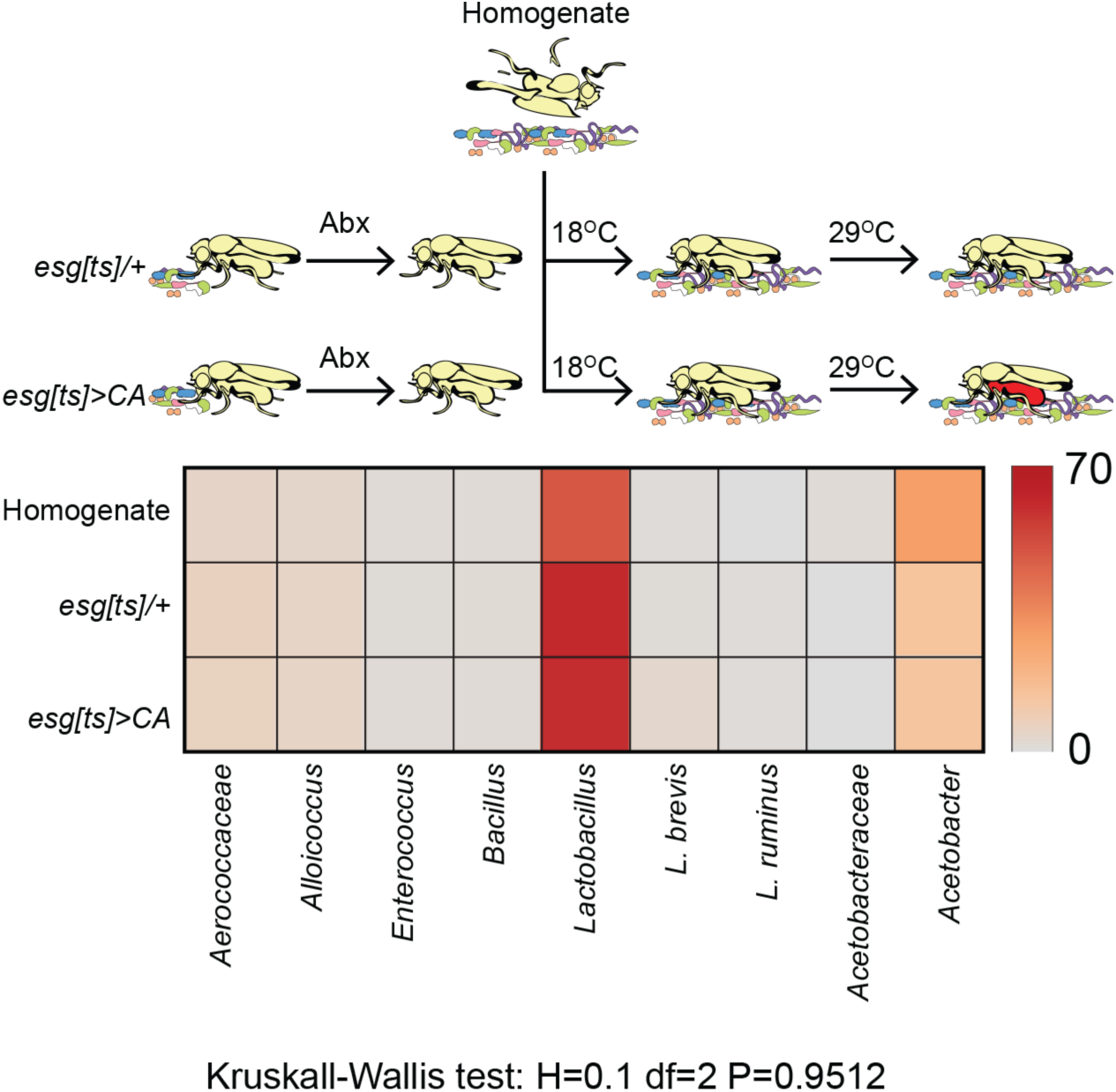
Induced expression of ImdCA in intestinal progenitor cells does not have a substantial impact on gut microbiome composition. Heat map analysis of microbial original taxonomic unit representation in dissected midguts of the indicated genotypes. All flies were raised under identical conditions, and fed the same homogenate at the same time. Kruskall-Wallis test results for degree of difference between the different samples.

### Constitutive IMD Pathway Activation Causes Dysplasia in Intestinal Progenitor Cells

In contrast to the minimal effects on bacterial microbiome composition, we noticed that ImdCA affected expression of regulators of intestinal homeostasis. This included elements of Wnt, Ras, Insulin, JNK and JAK/STAT pathways (Figure 5B). This led us to ask if ImdCA impacts intestinal morphology. To address this question, we examined the posterior midguts of two, eleven and twenty-six day old *esg^ts^>CA* and control *esg^ts^* flies. In these lines, the *esg^ts^* regulatory system marks all progenitors with GFP. We found that acute activation of the IMD pathway had no effect on midgut architecture. The posterior midguts of two-day old *esg^ts^* and *esg^ts^>CA* flies raised at 29°C were essentially indistinguishable (Figure 6A). In both cases, anti-Armadillo (beta-catenin ortholog) immunofluorescence showed a regular arrangement of large, polyploid ECs, that were interspersed with small, Prospero positive EE cells. Likewise, the posterior midguts of both genotypes contained regularly spaced nuclei and evenly distributed GFP positive progenitor cells (Figure 6A).

**Figure 5.**
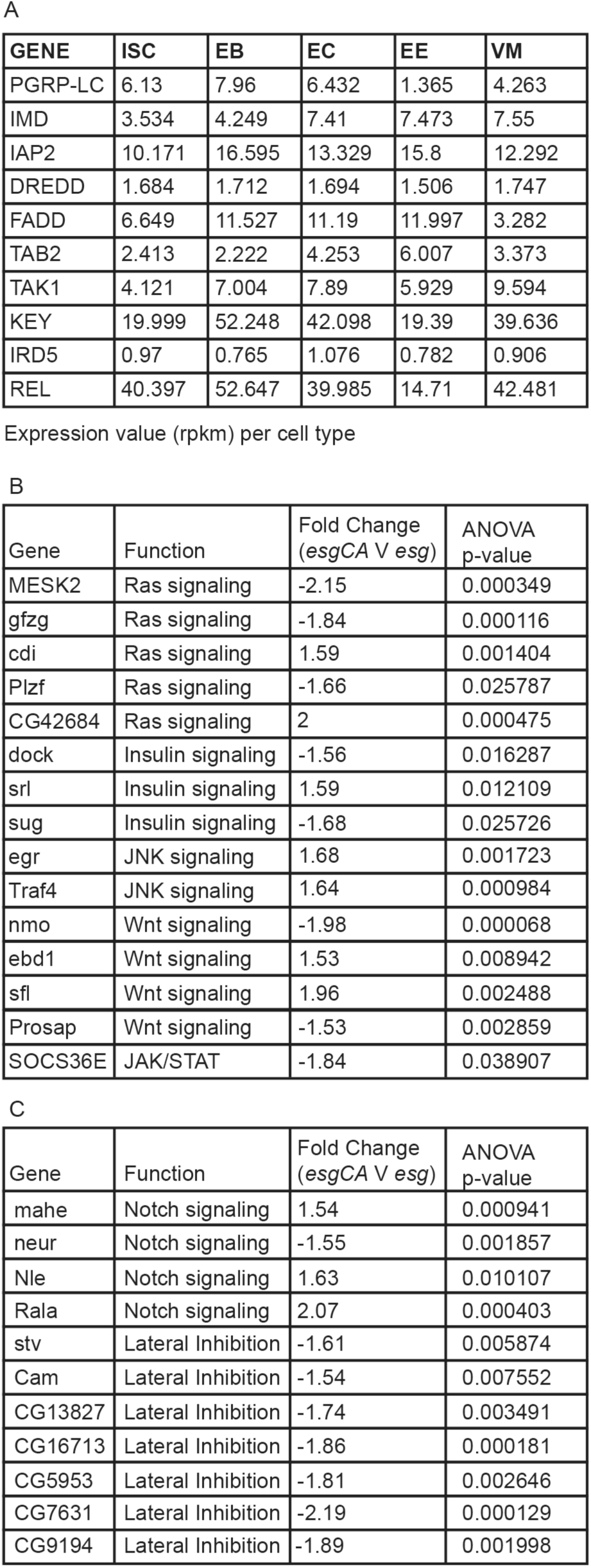
**A:** Expression of IMD pathway components in intestinal cell types. Expression data taken from (Dutta et al., 2015). **B-C:** Homeostatic regulators (B) and Notch pathway components (C) that were dysregulated in *esg^ts^>CA* flies relative to *esg^ts^* flies.

By day 11, the posterior midguts of *esg^ts^>CA* flies appeared moderately dysplastic relative to *esg^ts^* flies. By 26 days, the posterior midguts of control *esg^ts^* flies displayed classical hallmarks of age-dependent dysplasia such as a disorganized epithelium, uneven arrangements of nuclei, and irregular patterns of GFP expression (Figure 6A). We noticed that midgut architecture was further disrupted in 26 day-old *esg^ts^>CA* flies. In this case, we detected a near complete breakdown of epithelial organization (Figure 6A), an increase in cell density, and an increase in GFP positive cells. The hyperplastic phenotype of 26 day old *esg^ts^>CA* flies is not a consequence of accelerated physiological aging, as *esg^ts^* and *esg^ts^>CA* flies have indistinguishable lifespans (Figure 6I).

**Figure 6.**
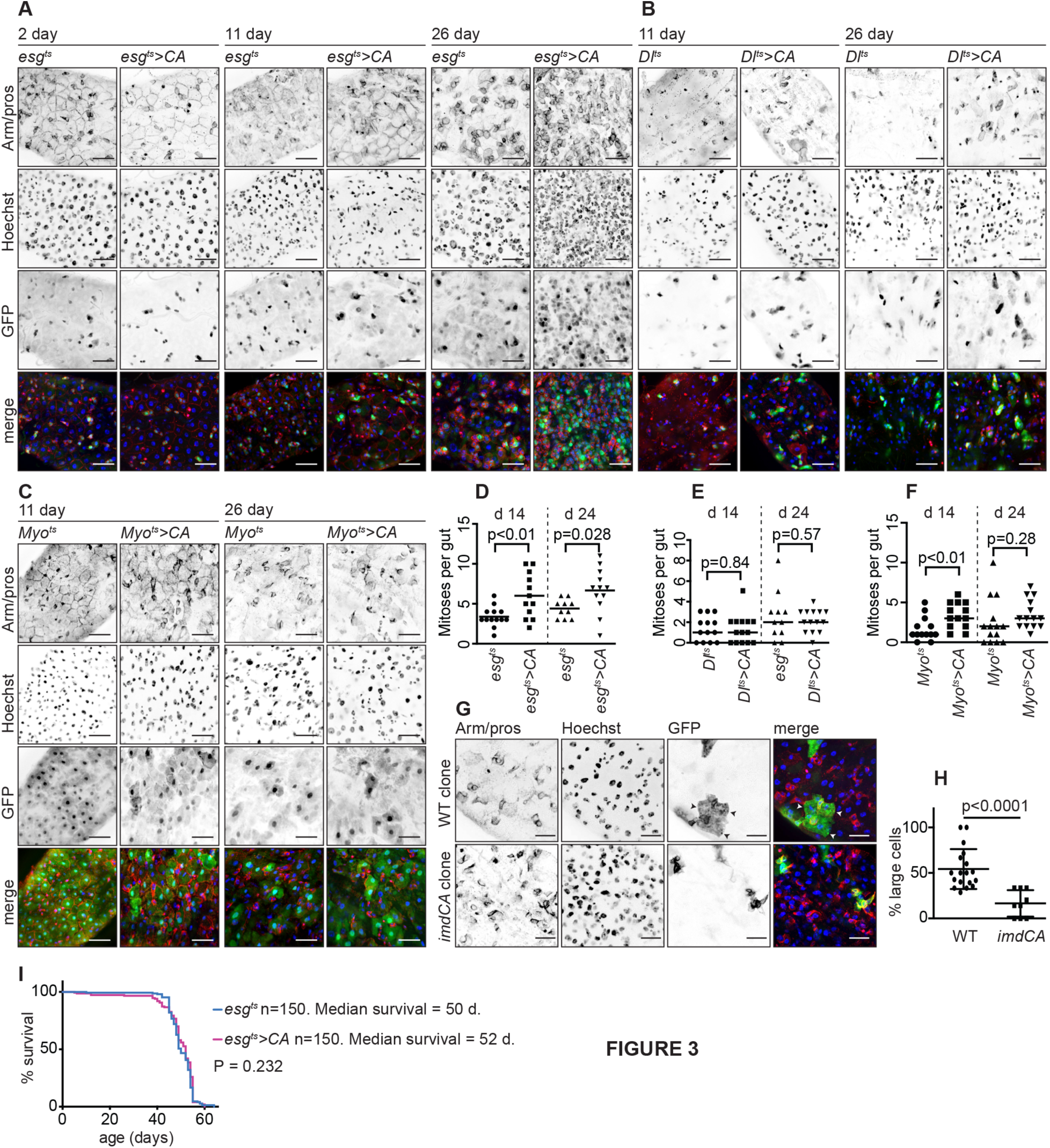
Intestinal hyperplasia in the posterior midguts of *esg^ts^>CA* flies. **(A-C)** Immunofluorescence imaging of the posterior midguts of *esg^ts^>CA* and *esg^ts^* flies (A); *Dl^ts^>CA* and *Dl^ts^* flies (B); and *myo1a^ts^>CA* and *myo1a^ts^* flies (C) raised at 29°C for 2, 11 or 26 days as indicated. Each midgut was stained with Armadillo (to indicate cell borders), and Prospero to reveal enteroendocrine cells; Hoechst to reveal nuclei; and GFP was visualized to reveal progenitor cells (A), stem cells (B) and enterocytes (C). All images were taken at 40X magnification and scale bars show 25 μm. **(D-F)** Number of mitotic cells in the intestines of flies of the indicated genotypes raised at 29°C for 14 and 24 days. **G:** Visualization of mitotic clones in wild type flies (upper row) or mitotic clones that express *imdCA* (lower row). All images were taken at 60X magnification and scale bars show 15 μm **H:** Percentage of large, enterocyte-type cells in wild type clones and *imdCA* positive clones. Comparisons were performed with a student’s t test. **(I)** Survival curves for 150 *esg^ts^>CA* and 150 *esg^ts^* flies raised at 29°C.

Given the hyperplasia and altered expression of homeostatic regulators in older *esg^ts^>CA* flies, we reasoned that ImdCA affects proliferation rates in intestinal progenitor cells. To test this hypothesis, we used anti-phosphorylated histone H3 immunofluorescence to count mitotic cells in the intestines of fourteen and twenty-four day-old *esg^ts^* and *esg^ts^>CA* files. At both ages, we detected a significant increase in mitotic events in *esg^ts^>CA* intestines relative to age-matched *esg^ts^* controls (Figure 6D). These data show that persistent expression of ImdCA in adult midgut progenitor cells disrupts expression of homeostatic regulators, increases cell proliferation, and promotes tissue hyperplasia.

We did not see a similar phenotype when we used a *Dl^ts^* line (*UASGFP*; *DlGAL4*, *GAL80^ts^*) to express ImdCA in GFP-marked stem cells. In this case, midgut morphology and proliferation rates were indistinguishable between *Dl^ts^>CA* and control *Dl^ts^/+* flies at all times (Figure 6B and E). Expression of ImdCA in GFP-marked intestinal enterocytes with the *Myo1A^ts^* (*UASGFP*; *Myo1aGAL4, GAL80^ts^*) line led to a mild increase in midgut proliferation at d14 that did not persist at later times (Figure 6C and F). Thus, expression of ImdCA in progenitor cells led to a marked hyperplasia throught the posterior midgut. As the *Dl^ts^* line, fails to induce hyperplasia our data suggest that activation of IMD either in progenitor pairs, or enteroblasts alone leads to midgut hyperplasia.

### Persistent IMD Signals Disrupt Notch Activity

Among homeostatic regulators, ImdCA had a pronounced effect on expression of Notch pathway components (Figure 5B). This observation matches an earlier report of deregulated expression of Notch pathway genes in *imd* mutant flies (Broderick et al., 2014), and raises the possibility that the IMD pathway influences Notch activity in the adult. To test if ImdCA affects progenitor cell differentiation, we used MARCM analysis to identify individual mitoses and subsequent differentiation events in adult posterior midguts. As expected, mitotic clones in conventional flies contained a mix of smaller cells (most likely progenitors and EE), and larger, polyploid ECs (Figure 6G). In contrast, we rarely detected large, polyploid cells in mitotic clones that express ImdCA. Instead, ImdCA clones typically contained aggregates of small cells (Figure 6G). These observations suggest that persistent expression of ImdCA in progenitor cells affects the differentiation of EBs into ECs.

To directly test the possibility that ImdCA disturbs EB differentiation, we determined the percentage of EE cells in the intestines of twenty-four days old *esg^ts^* and *esg^ts^>CA* flies. As expected, the posterior midguts of *esg^ts^>CA* flies contained greater numbers of cells per area than those observed in age-matched, control *esg^ts^* flies (Figure 7A-C). However, we also found that a significantly greater percentage of midgut cells expressed the EE marker Prospero in *esg^ts^>CA* than in *esg^ts^* midguts (Figure 7A, B, D). These data show that persistent expression of ImdCA disrupts the developmental trajectory of progenitor cells, and leads to the differentiation of supernumerary EE.

**Figure 7.**
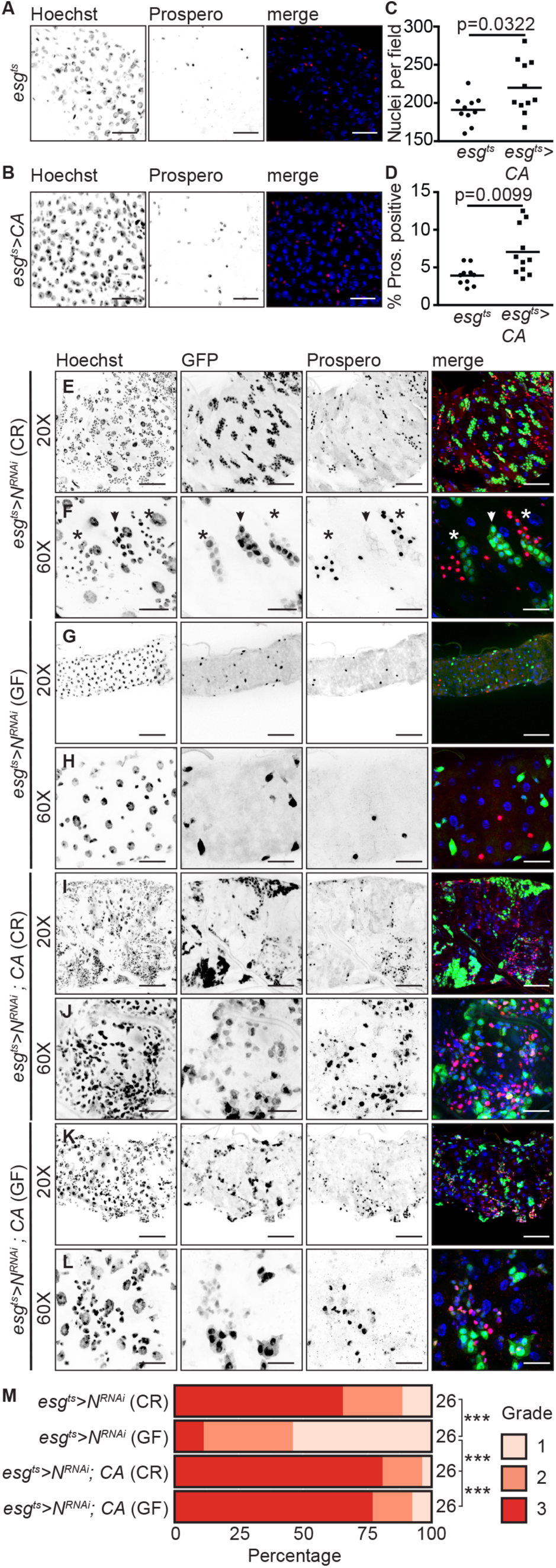
ImdCA promotes tumorigenesis in Notch-deficient progenitor cells. **(A-B)** Midguts of 20d old *esg^ts^* (A) and *esg^ts^>CA* (B) flies stained for DNA and Prospero as indicated. Images were false-colored and merged with DNA in blue and Prospero in red. Images were taken at 40X magnification and scale bars show 25 μm. **C:** Automated quantification of total nuclei in a given field for flies of both genotypes. **D:** Total percentage of Prospero positive cells in the same fields. For C and D comparisons were performed with a student’s t test. **E-L:** Visualization of tumorigenesis in 8d old *esg^ts^>N^RNAi^* and *esg^ts^>N^RNAi^*; *CA* flies. Posterior midguts were stained for Prospero and DNA in conventionally-reared *esg^ts^>N^RNAi^*; flies (E-F); germ-free *esg^ts^>N^RNAi^* flies (G-H); conventionally-reared *esg^ts^>N^RNAi^*; *CA* flies (I-J); and germ-free *esg^ts^>N^RNAi^*; *CA* flies (K-L). In all cases, progenitor cells were visualized through the expression of GFP. Scale bars show 50μm for 20X magnification images, and 15μm for 60X images. A representative grade 2 tumor is indicated with an arrowhead in F, and representative grade 3 tumors are shown with an asterisk. **M:** Tumor scoring in the posterior midguts of 8d old flies with the indicated genotypes and culture conditions (n=26) for each condition. Tumor grade was compared between individual treatments using one-way ANOVA, with Tukey’s correction. indicates P<0.0001.

### IMD Pathway Activity Promotes the Development of Notch-Mediated Intestinal Tumors

As bacterial challenges enhance *Notch*-deficient tumor formation (Apidianakis et al., 2009, Jiang et al., 2009), and ImdCA influences *Notch*-dependent events, we asked what effects persistent immune signals have on models of *Notch*-deficient tumorigenesis in the posterior midgut. To answer this question, we used a validated, inducible RNAi line that blocks *Notch* signaling in GFP-marked adult progenitors (*esg^ts^>N^RNAi^*). As anticipated, the posterior midguts of *esg^ts^>N^RNAi^* flies contained GFP-positive tumors composed of small, irregularly spaced nuclei within eight days of Notch depletion (Figure 7E, F). Many tumors also contained patches of Prospero-positive enteroendocrine cells (Figure 7F). The ability of *esg^ts^>N^RNAi^* to promote tumorigenesis appears to require extrinsic inputs from the microbiome, as removal of the gut microbiome greatly reduced the incidence of tumor formation in *esg^ts^>N^RNAi^* posterior midguts (Figure 7G, H). This observation matches an earlier report that growth of *Notch* loss-of-function tumors requires stress-induced mitogenic cues (Patel et al., 2015).

Whereas elimination of the microbiome blocked tumorigenesis in *esg^ts^>N^RNAi^* flies, co-expression of ImdCA with *^NRNAi^* (*esg^ts^>N^RNAi^*; *CA*) exacerbated the formation of midgut tumors. We invariably detected large patches of GFP and Prospero positive cells in *esg^ts^>N^RNAi^*; *CA* posterior midguts, with few large cells characteristic of mature ECs (Figure 7 I, J). To determine if ImdCA alone provides sufficient stress signals to promote tumorigenesis in this *Notch* loss-of-function model, we examined the posterior midguts of germ-free *esg^ts^>N^RNAi^*; *CA* flies. We found that nearly all midguts of germ-free *esgts>CA, ^NRNAi^* flies had accumulations of small, GFP-positive cells, large patches of Prospero positive tumors, and few large polyploid ECs (Figure 7K, L). To quantify the effects of microbiome elimination on tumorigenesis, we developed a grading system for tumor scale in the posterior midgut. We defined tumor-free intestines as Grade 1 (e.g. Figure 7G, H), tumors that only contained GFP-positive clusters of cells as Grade 2 (Figure 7F, asterisks), and tumors that contained clusters of GFP-positive and Prospero-positive cells as Grade 3 (Figure 7F, arrowhead). We used this system to quantify tumor grade in 26 intestines from each experimental condition. This scale confirmed a significant reduction of tumor formation in germ-free *esg^ts^>N^RNAi^* flies (Figure 7M), and a significant increase of tumorigenesis in germ-free *esgts>N^RNA^; CA* flies (Figure 7M).This phenotype is remarkably similar to one described earlier for bacterial infection of *esg^ts^>N^RNAi^* flies (Apidianakis et al., 2009, Jiang et al., 2009), and shows that immune signaling in progenitor cells alone is sufficient to promote *Notch*-dependent tumor growth in adult midguts.

## DISCUSSION

Animal genomes encode evolutionarily conserved systems that mitigate the threats posed by potentially harmful microbes. For example, homeostatic resistance systems clear invading microbes, while tolerance mechanisms limit the damage caused by invaders (Medzhitov et al., 2012, Schneider and Ayres, 2008). In this study, we used the genetically accessible fruit fly to activate immune responses exclusively in intestinal progenitor cells. We focused on progenitors, as they are immune-competent, but comparatively understudied from the perspective of IMD. We showed that persistent immune activity causes intestinal hyperplasia, disrupts progenitor cell differentiation, and fuels Notch-dependent tumorigenesis. These phenotypes occur without significant shift in bacterial populations, consistent with altered disease tolerance in the host. Our studies match reports of relationships between the microbiota, inflammation and colorectal cancer (Belkaid and Hand, 2014, Irrazabal et al., 2014), and uncover a novel requirement for regulated innate immune activity in the overall maintenance of progenitor cell homeostasis in the fly.

In *Drosophila*, several homeostatic devices control progenitor cell proliferation and differentiation, often in response to bacterial products (Amcheslavsky et al., 2009, Buchon et al., 2009, Jiang et al., 2009). For example, flies raised in the absence of a microbiome undergo fewer mitoses in the midgut (Buchon et al., 2009). In addition, flies raised in a germ-free environment have atypical numbers of prospero-positive EE cells, a lower density of ECs, and deregulated expression of Notch pathway components (Broderick et al., 2014). It is unclear how much of this response requires an intact IMD pathway, although recent transcriptional work established that mutations of the IMD pathway impact numerous aspects of intestinal transcription (Broderick et al., 2014, Erkosar et al., 2014), including Notch pathway components (Broderick et al., 2014). Our work shows that expression of ImdCA in progenitor cells alone recapitulates key features of those host responses to gut microbes. For example, of the 253 genes differentially expressed in the intestines of conventionally-reared flies compared to germ-free flies, 87 are affected by expression of ImdCA. Additionally, expression of ImdCA in progenitor cells boosts intestinal mitoses, affects EE cell differentiation, and increases EC density. Combined, these data implicate the IMD pathway as a novel regulator of progenitor cell proliferation.

As the microbiome induces progenitor cell proliferation, we initially assumed that ImdCA-dependent hyperplasia was the result of a dysbiosis in the gut caused by immune activity. However, expression of ImdCA does not influence microbiome composition. These data raise the possibility that common intestinal microbes are insensitive to IMD activity in the gut, or that elevated immune signaling activates counter-defenses in commensal populations, thereby allowing persistence of commensal bacteria. This matches experimental data that commensal bacteria effectively overcome host innate defenses in other models. For example, many gut-resident *Firmicutes* and *Bacteroidetes* are not susceptible to antimicrobial peptides (Cullen et al., 2015), and NOD deficient mice do not display dysbiosis (Robertson et al., 2013, Shanahan et al., 2014). These observations lead us propose that constitutive activation of IMD acts directly on the host to induces intestinal hyperplasia. Although out data are more consistent with IMD acting in enteroblasts to promote hyperplasia, additional experiments are required to rule out a need for IMD activity in progenitor pairs. We believe that the ability of ImdCA to promote Notch-dependent tumors is of particular significance in the context of aging. As flies age, clones of *Notch* mutant cells spontaneously develop in the intestine (Siudeja et al., 2015), and host immune activity gradually rises (Buchon et al., 2009). Our data suggest that age-dependent increases in immune activity may contribute to the formation, or growth, of Notch-dependent tumors in the fly. Indeed, we found that elimination of the microbiome was sufficient to arrest the appearance of Notch-dependent tumors.

The role of Notch in tumorigenesis is not restricted to flies. Several studies established clear links between deregulated Notch activity and carcinogenesis in vertebrates (Rizzo et al., 2008, Shih Ie and Wang, 2007). Colorectal cancer lines and adenocarcinomas express elevated levels of Notch1 and Notch 2 (Fre et al., 2009, Guilmeau et al., 2010, Reedijk et al., 2008, Sikandar et al., 2010). In addition, manipulations of Notch pathway activity modify tumorigenesis in mouse models of colorectal cancer (Fre et al., 2009, Rodilla et al., 2009). The mechanisms that activate Notch in colon cancer are unclear at present. Given the broad conservation of immune activity and Notch signaling in the guts of flies and mammals, our work raises the possibility that chronic inflammation is a long-term activator of Notch-dependent tumorigenesis. The data in this study present a simple model to precisely describe the relationships between intestinal immune activity and the formation of Notch-dependent tumors.

## EXPERIMENTAL PROCEDURES

### *Drosophila* husbandry

All flies were raised on standard corn meal medium (Nutri-Fly Bloomington Formulation, Genesse Scientific). Germ-free husbandry conditions are described in the Supplementary methods. For 16S deep-sequencing, we raised freshly eclosed virgin females flies on antibiotic medium for 5 d at 18°C, then switched to sterile antibiotic-free food for 1 d. Next, flies were fed a homogenate prepared from *w^1118^* flies for 16 h. Afterwards, flies were raised on sterile antibiotic-free food at 18°C for 7 d. Flies were moved to 29°C for another 7 days, and 10 guts/sample were dissected. We extracted microbial DNA with the UltraClean^®^ Microbial DNA Isolation Kit (MoBio Laboratories Inc.), and amplified 16S DNA with Platinum^®^ PFX Taq (Invitrogen), followed by purification with the QIAquick^®^ PCR Purification Kit (Qiagen). Concentration was measured on the Qubit^®^ 2.0 (Invitrogen) and 1 ng was used for library prep. Libraries were prepared using the Nextera XT DNA Library Preparation Kit (Illumina). We purified libraries using Ampure Beads (Qiagen) and quantified using the Qubit^®^ 2.0 (Invitrogen) and Bioanalyzer 2100 (Agilent). Pooled libraries were loaded on the Miseq (illumina) using the MiSeq Reagent Kit v3 (600-cycle) for sequencing. 16S sequences were assembled using DNASTAR Navigator, and annotated with the greengenes database. For infection studies, freshly eclosed flies were raised for 5 d at 18°C, then switched to 29°C for 2 d. 24 h before infection 100 μl of a *V. cholerae* C6706 glycerol stock was spread on LB agar plates and grown at 29°C. Flies were starved for 2 h before infection. The bacterial lawn was scraped from the plate and mixed into LB broth, then diluted to an OD600 of 0.125, flies were fed 3ml bacterial supernatant on a cotton plug, and dead flies counted every 2-4 h. To generate mitotic clones, flies were raised at 18°C for 5-6 d, incubated at 37°C for 2 h, and raised for an additional 8-10 days at 25°C.

### Cell culture and molecular biology

The Imd expression construct used in this study has been described elsewhere (Guntermann and Foley, 2011). To generate ImdCA, we used pENTR/D-TOPO ImdCA forward and reverse primers (supplemental material) to amplify a truncated Imd. We then cloned truncated Imd with into the pENTR/D-TOPO plasmid according to manufacturer’s instructions (Invitrogen). We recombined TOPO-ImdCA with pAWM, or pTW (LR recombination, Invitrogen) to generate ImdCA-6myc and UAS-ImdCA expression plasmids. Transgenic lines were generated by Bestgene Inc. Immunoprecipitation and Western blot protocols have been described elsewhere (Guntermann and Foley, 2011). For immunoprecipitation assays, we incubated cell lysate with mouse anti-HA (Sigma, 1:500) at 4°C overnight, added protein G-Sepharose beads and incubated for an additional hour at 4°C. Beads were pelleted by centrifugation at 300 X g for 30 s and washed in lysis buffer three times. After discarding the supernatant, beads were re-suspended in 2X sample buffer, and analyzed by Western blot.

### Gene expression analysis

We used TRIZOL to prepare RNA from dissected adult intestines (Guntermann and Foley, 2011). Microarray studies were performed in triplicate on virgin flies that we raised on regular or antibiotic-treated food food for 10-11 days at 29°C. We then shifted flies to 29°C for another 2 days, after which guts were dissected for RNA extraction (5 females and 5 males per sample). We used 100 ng purified RNA to make labeled cRNA for microarrays using the GeneChip^®^ 3’ IVT Plus Reagent Kit (Affymetrix), then fragmented cRNA and then hybridized to the GeneChip^®^Drosophila Genome 2.0 Array (Affymetrix). Preliminary analysis was done in the Transciptome Analysis Console (TAC) software (Affymetrix). We analyzed gene expression data using FlyMINE (Lyne et al., 2007) and Panther (Thomas et al., 2003). Array data has been submitted to the NCBI GEO database (accession ID: GSE89445). For qPCR studies, we prepared 1 μg cDNA from purified RNA using qScript cDNA Supermix (Quanta Biosciences, Inc.) and then performed qPCR using the PerfeCTa^®^ SYBR^®^Green FastMix (Quanta Biosciences, Inc.). All qPCR studies were performed in triplicate and relative expression values were calculated using delta delta Ct calculations. In each case, expression levels were normalized to *actin.*

### Immunofluorescence

The immunofluorescence protocol used in this study has been described elsewhere (Petkau et al., 2014). All immunofluorescence images were prepared from the posterior midgut of adult flies at a distance of approximately 50 μM from the hindgut transition. To quantify phospho-histone H3 positive cells, guts were visualized under the microscope and scanned from posterior midgut (hindgut transition area) to anterior midgut (crop). To quantify Prospero positive cells, we used the Columbus software to identify the percentage of Hoechst positive cells that were also prospero positive. 3-D reconstructions were created with Volocity^®^ 6.3 (Perkin Elmer).

### Germ-Free Adult Fly Husbandry

Larval growth and development is supported by commensal microbes. As our primary goal was to examine the effects of immune signaling in an adult model of intestinal tumorigenesis, we were concerned that elimination of the larval microbiome would influence gut development, and possibly confound interpretation of our data. For that reason, we chose to raise larvae under conventional conditions, and exclusively remove the bacterial microbiome from adults. To generate germ-free animals, we raised freshly eclosed adult flies on sterile food containing an antibiotic cocktail (100 μg/ml Ampicillin,100 μg/ml Neomycin, 100 μg/ml Metronidazole and 50 μg/ml Vancomycin). This protocol allowed us to raise larvae in the presence of a conventional microbiome, and restrict our examination of germ-free phenotypes to the adult life stage. We confirmed bacterial elimination by plating fly homogenates on agar plates permissive for the growth of *Lactobacilli* and *Actetobacter.*

### Primers used in this study

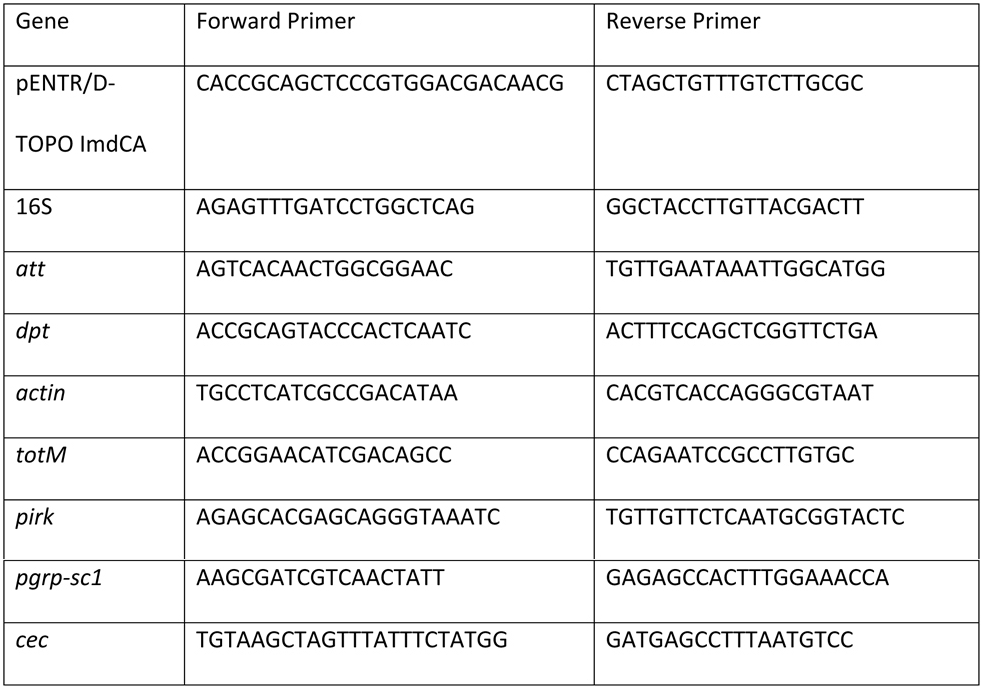

### Antibodies used in this study

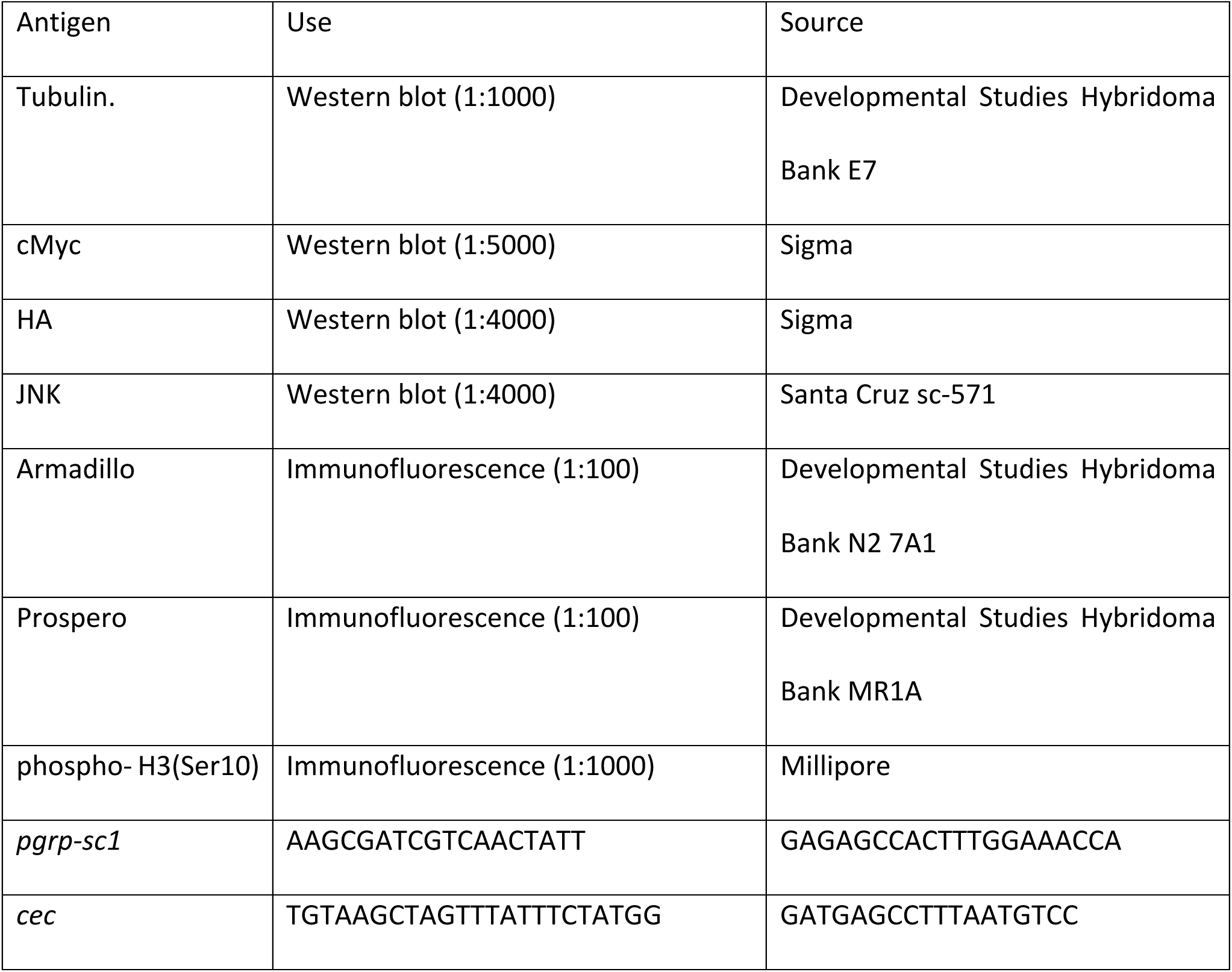

### Fly Genotypes

Clonal analysis was performed with flies of the following genotypes: *hsFLP, UASGFP/X*; *tubGAL80, neoFRT40A/neoFRT40A*; *tubGAL4/+* (wild-type), and *hsFLP, UASGFP/X*; *tubGAL80, neoFRT40A/neoFRT40A*; *tubGAL4/UASimdCA* (ImdCA positive clones).

## AUTHOR CONTRIBUTIONS

K.P., M.F., S.G., and E.F. conceived and designed experiments; K.P., M.F., and S.G. performed the experiments; K.P., M.F., and E.F performed data analysis and wrote the paper.

## ACKNOWLEDGEMENTS

Transgenic flies were provided by Bruce Edgar (*esg^ts^*), Bexacerbated the formationruno Lemaitre (*dredd*), the Bloomington *Drosophila* stock Center (*cgGAL4* and *GAL80^ts^*), and the Vienna *Drosophila* Resource Center (*N^RNAi^*).The research was funded by a grant from the Canadian Institutes of Health Research to EF (MOP 77746). We acknowledge the microscopy support from Dr. Stephen Ogg and the Faculty of Medicine and Dentistry core imaging service, the Cell Imaging Centre, University of Alberta. Microarrays were processed at the Alberta Transplant Applied Genomics Center. We are grateful to Juan Jovel for bioinformatics advice.

